# Visualizing phenotypic heterogeneity and single-cell morphology *in situ* during gut infection

**DOI:** 10.1101/2025.04.30.651596

**Authors:** Nicholas V. DiBenedetto, M. Lauren Donnelly-Morell, Carol A. Kumamoto, Aimee Shen

## Abstract

Visualizing phenotypic heterogeneity at single-cell resolution within dense microbial communities is technically challenging. Here, we present a method for visualizing this heterogeneity by combining spectrally compatible reporters to track the spatial distribution of gene expression in individual bacterial cells in the mammalian gut. Using toxin gene expression in *Clostridioides difficile* as a model for visualizing phenotypic heterogeneity, we demonstrate that, while *C. difficile* primarily occupies the lumen, a subpopulation of *C. difficile* associates with the colonic epithelium independent of toxin production. The approach further revealed that heterogeneity in *C. difficile* toxin gene expression is independent of location in the gut and unexpectedly showed that a toxin gene over-expressing mutant forms filamentous cells during the acute phase of infection. Thus, our reporter system provides quantitative, single-cell resolution of bacterial behavior within the intact gut environment and establishes a broadly applicable platform for investigating phenotypic heterogeneity in dense microbial communities.

## Introduction

Localizing bacteria in their native context within a host has provided critical insights into host-microbe interactions and bacterial behavior in complex environments^1–5^. While methods such as fluorescence *in situ* hybridization (FISH) have visualized the spatial organization of bacterial species within tissues^6,7^, these analyses do not provide information on the locations of phenotypically distinct sub-populations of a given species. Notably, fluorescent reporter strains have allowed phenotypic variation at the single-cell level in bacteria to be visualized *in situ* with minimal handling relative to techniques like RNA-FISH^8^. For example, *Vibrio cholerae* upregulates toxin genes during biofilm formation in the small intestine^9^, *Yersinia pseudotuberculosis* spatially regulates virulence and detoxification gene expression in deep tissue sites^10,11^, and *Mycobacterium tuberculosis* exhibits differential replication rates depending on its location within a caseous lesion^12,13^. While these analyses have revealed how specific bacterial sub-populations adapt to and interact with distinct microenvironments, the tissues infected by these bacteria have considerably lower microbial complexity than that found in the colon.

Historically, localizing bacteria *in situ* in the colon has been challenging due to the high density and dynamic nature of bacterial populations within this site^14,15^. While most FISH analyses have not localized specific bacterial species within the gut^7,16^, fluorescent reporters have achieved visualization of individual *Bacteroides* spp. at the single-cell level *in situ* during colonization^17,18^. However, this work was done either in germ-free or weaning mice, which does not fully model the microbial complexity or immunocompetency of a conventional mouse model. Other studies have used fluorescent reporters to localize colonic pathogens like *Shigella sonnei* during infection, but these analyses have used non-natural infection routes and inocula^19^ to facilitate the visualization of pathogens *in situ*.

Developing methods for visualizing phenotypic heterogeneity *in situ* in the colon would significantly advance our understanding of how specific bacterial species adapt to this densely populated and competitive environment^20^. Indeed, immense selective pressures promote the generation of phenotypically heterogeneous sub-populations^21^. For example, bacterial capsule genes in *Bacteroides*^22^ and flagella and Type III secretion system genes in *Salmonella*^23,24^ are heterogeneously expressed to promote colonization and virulence, respectively. Unfortunately, it has been technically challenging to reliably visualize phenotypic heterogeneity *in situ* in the colon, leaving questions about the frequency, spatial distribution, and selection of specific phenotypes during infection.

Visualizing phenotypic heterogeneity in the major nosocomial pathogen, *Clostridioides difficile*, is particularly relevant because numerous traits, such as flagellar motility, cell chaining, and sporulation, are heterogeneously expressed by this pathogen^21,25,26^. In addition, *C. difficile* expresses toxin genes in a bimodal manner *in vitro*^27^, although the dynamics of toxin gene expression at the single-cell level during infection remain unclear. Understanding the location, frequency, and magnitude of toxin gene expression during infection is critical because toxin production is essential for disease^28^, and the ability of toxins to bind their receptors on the colonic epithelium likely impacts disease severity^29^. In addition, toxin production is a critical factor for diagnosing *C. difficile* infection^30,31^, but the location of its production could impact the progression of the disease.

To address these challenges, we developed fluorescent reporter strains for visualizing toxin gene expression at the single-cell level during the natural infection of conventional mice by *C. difficile*. We found that toxin gene expression is heterogeneously distributed *in situ* and that *C. difficile* is found in diverse environments in colonic sections, including in close association with the epithelial cell layer, a spatial localization that has not been previously described for *C. difficile* in conventional mice. Moreover, we discovered that the relative localization of *C. difficile* within the colon does not strongly impact the frequency or magnitude of toxin gene expression and that toxin production did not affect where *C. difficile* localizes in the colon. Remarkably, these imaging analyses revealed that a toxin gene overexpressing mutant forms filamentous cells during the acute phase of infection. Thus, our methodology provides a robust and streamlined workflow for visualizing phenotypic heterogeneity in *C. difficile* that can likely be applied to other clostridial species. Our analyses further highlight how visualizing bacteria *in situ* can reveal previously unknown morphologies that are not captured *in vitro*, offering new insights into microbial behavior and disease pathogenesis.

## Results

### Optimization of constitutive fluorescent reporters for visualizing *Clostridioides difficile*

To visualize the subset of *Clostridioides difficile* cells that express toxin genes during infection, we first developed spectrally compatible, constitutive fluorescent reporters that allow individual *C. difficile* cells to be visualized within a colonic section. Building upon chromosomally-encoded constitutive *mNeonGreen* (*mNG*) reporters previously developed for *C. difficile*^32^, we first sought to reduce the amount of mNG produced by the P*_slpA_::LP-mNG* strain because its high-level mNG production caused toxicity at late stages of growth *in vitro*. To this end, we deleted the sequence encoding a leader peptide (*LP*) (Fig. 1a) previously shown to enhance the fluorescence of reporter constructs in *Bacteroides* spp.^17^. Loss of the LP reduced P*_slpA_::mNG* reporter fluorescence by 8-fold (Fig. 1b,c). We also tested a P*_gluD_::mNG* reporter construct because RNA-Seq analyses indicated that this promoter is highly expressed during murine infection^33^. Indeed, *gluD* encodes a glutamine dehydrogenase that is used in diagnostic tests for detecting *C. difficile* infections^31^. The P*_gluD_::mNG* reporter was 3-fold dimmer than the *P_slpA_::mNG* reporter developed here, and it was also dimmer than the P*_cwp2_::LP-mNG* reporter we previously developed^32^ (Fig. 1b,c). Notably, aside from the P*_slpA_::LP-mNG* strain, the *mNG* reporter strains exhibited uniform fluorescence and WT growth during broth culture (Fig. 1b-d).

**Fig. 1:**
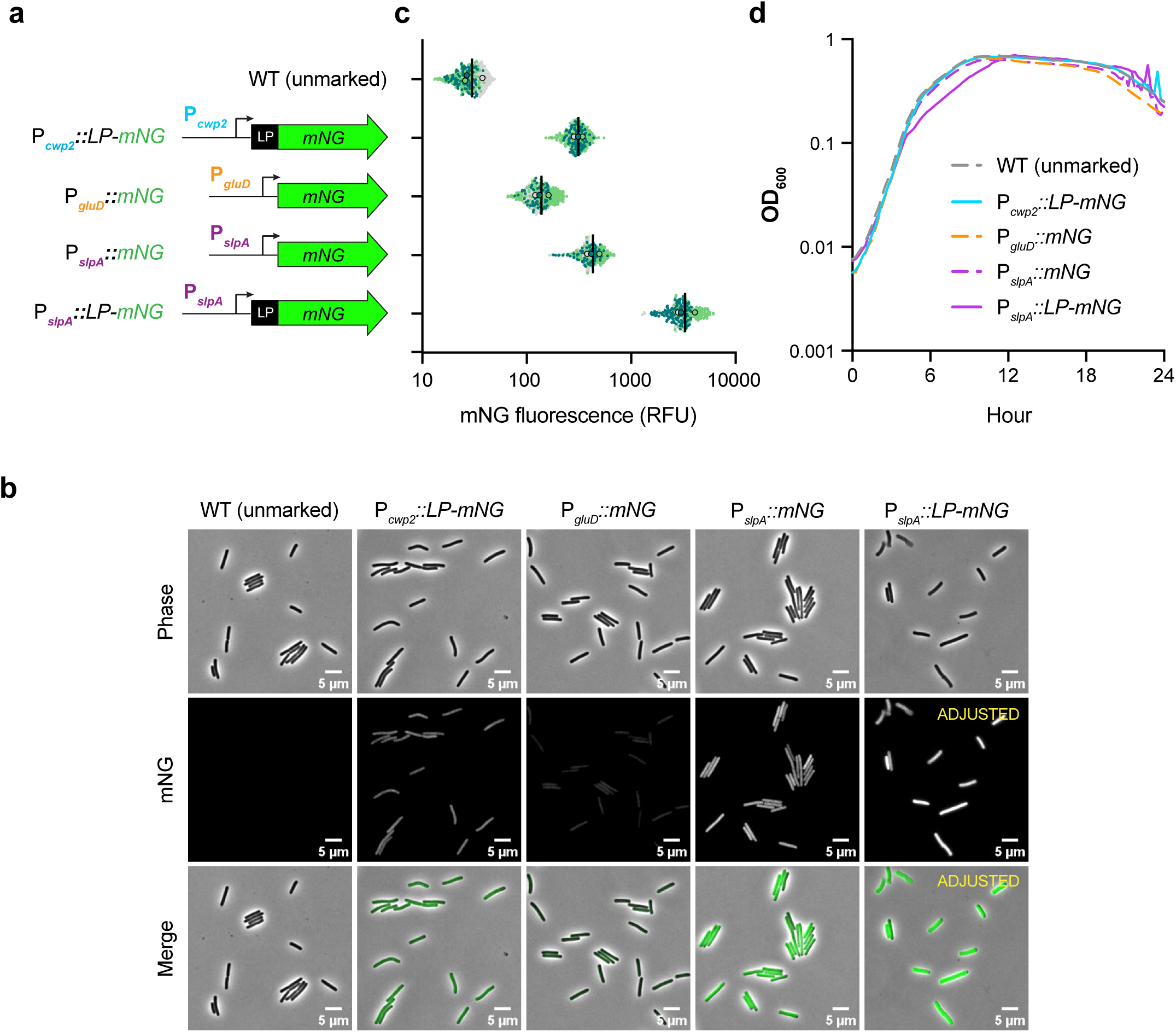
Optimization of constitutive *mNeonGreen* reporters. **a.** Schematic of constitutive reporter constructs with the indicated promoters driving expression of *mNeonGreen* (*mNG*). LP encodes a leader peptide (*LP*) that functions to increase mNG levels. **b.** Fluorescence microscopy of fixed cells expressing *mNG* grown to mid-log phase in BHIS broth culture. The mNeonGreen signal for P*_slpA_::mNG* was reduced in images marked “ADJUSTED” because the reporter strain is 8-24-fold brighter than the other *mNG* reporter strains. Scale bar is 5 µm. **c.** Superplots of single-cell fluorescence intensity quantified using SuperSegger^56^ for the strains shown in **b**. The outlined colored dots represent the median fluorescence measured for a given biological replicate. The horizontal black line indicates the mean fluorescence value determined for three biological replicates (unmarked WT n = 564; P*_cwp2_::LP-mNG* n = 499; P*_gluD_::mNG* n = 538; P*_slpA_::mNG* n = 459; P*_slpA_::LP-mNG* n = 327). **d.** Growth of the constitutive *mNG* reporter strains in BHIS broth based on optical density. A single biological replicate representative of three biological replicates is shown.

In parallel, we optimized spectrally compatible *mScarlet*-based constitutive reporters. While we previously generated P*_cwp2_::LP-mSc* and P*_slpA_::LP-mSc* reporter strains^32^, the recently developed mScarletI3 (mScI3) variant is ∼3-fold brighter and matures more rapidly than mScarlet^34^. Notably, replacing the *mSc* reporter with the *mScI3* reporter increased fluorescence by 6-fold for the P*_cwp2_* constructs (Fig 2a-c). In the course of cloning the P*_cwp2_::LP-mScI3* construct, we serendipitously determined that a G228R substitution enhanced the brightness of mScI3 in *C. difficile* by 2-fold (Fig. 2c). Expressing *mScI3* under the control of P*_slpA_* without the leader peptide sequence increased the mean fluorescence by ∼3-fold and led to more uniform fluorescence than the *mSc* reporters, likely because mScI3 matures faster than mSc^34^. Similar to our findings with the *mNG* reporter, the P*_gluD_::mScI3* construct was the dimmest of the three promoter constructs tested (Fig. 2b, c). Importantly, high-level *mSc* or *mScI3* expression had minimal impact on bacterial growth even for the P*_slpA_*-driven constructs (Fig. 2d).

**Fig. 2:**
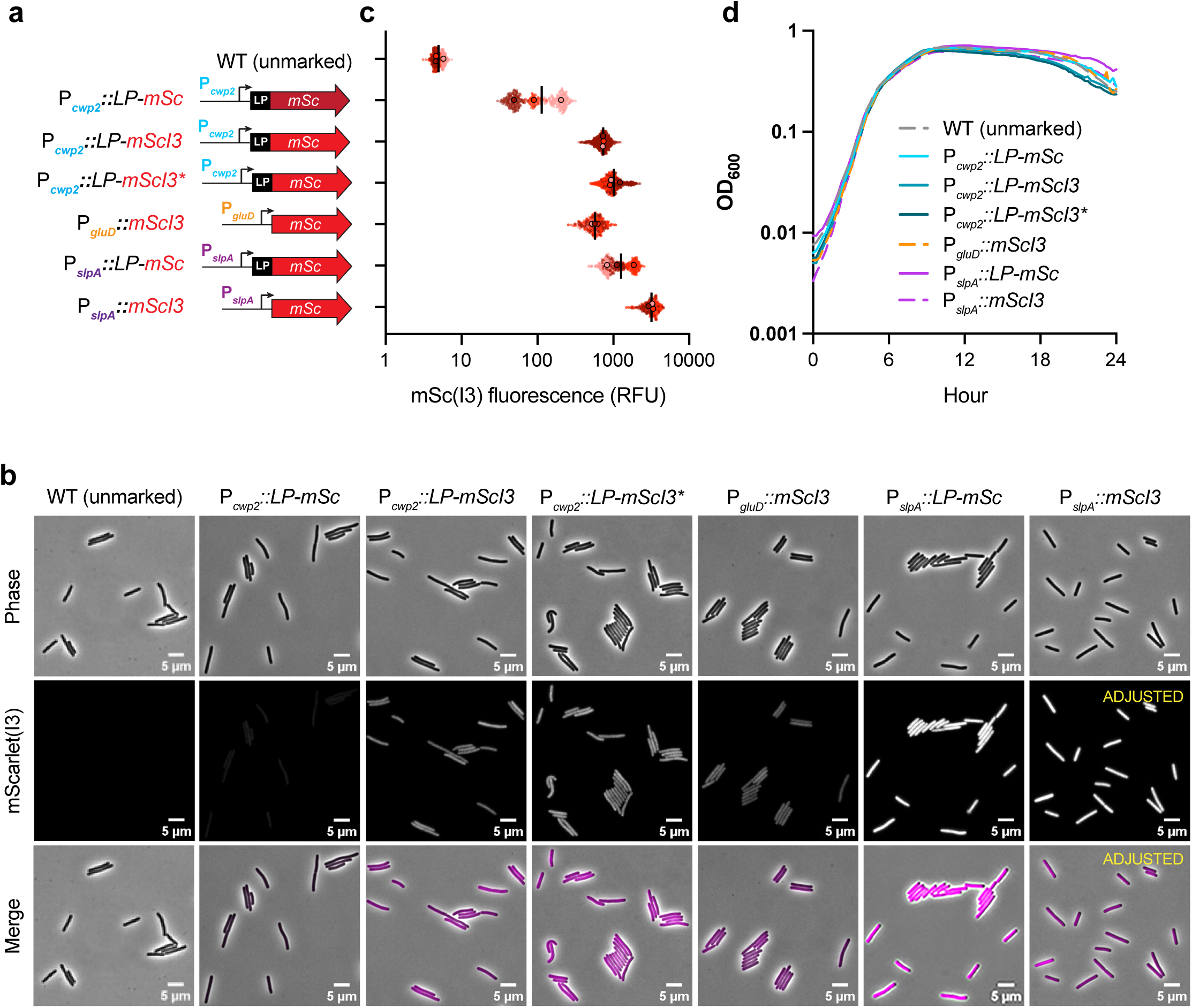
Optimization of constitutive *mScarlet*(*I3*) reporters. **a.** Schematic of constitutive reporter constructs with the indicated promoters driving expression of *mScarlet* (*mSc*) or *mScarletI3* (*mScI3*). *LP* encodes a leader peptide that functions to increase mSc(I3) levels. **b.** Fluorescence microscopy of fixed cells expressing *mScarlet* or *mScarletI3*. The mScI3 signal was reduced in images marked with “ADJUSTED” because the P*_slpA_::mScI3* reporter strain is 3-28-fold brighter than the other *mSc(I3)* reporter strains. Scale bar is 5 µm. **c.** Superplots of single-cell fluorescence intensity quantified using SuperSegger^56^ for the strains shown in **b**. The outlined colored dots represent the median fluorescence measured for a given biological replicate. The horizontal black line indicates the mean fluorescence value determined for three biological replicates (unmarked WT n = 547; P*_cwp2_::LP-mSc* n = 391; P*_cwp2_::LP-mScI3* n = 760; P*_cwp2_::LP-mScI3\** n = 743; P*_gluD_::mScI3* n =4 43; P*_slpA_::LP-mSc* n = 498; P*_slpA_::mScI3* n = 376). **d.** Growth of the constitutive *mSc*(*I3*) reporter strains in BHIS broth based on optical density. A single biological replicate representative of three biological replicates is shown.

### Visualization of constitutive *C. difficile* fluorescent reporter strains *in vivo*

Having identified *mNG* and *mScI3* reporter constructs that allow for stable, robust fluorescence *in vitro*, we next tested whether the constitutive reporter strains could be visualized during murine infection. Eight-week-old antibiotic-treated, female C57BL/6 mice were orally gavaged with 1 x 10^6^ *C. difficile* spores derived from reporter strains. Infected mice were sacrificed 48 hours post-infection, when peak disease severity is typically observed^35^. Colonic tissues were harvested, embedded in OCT, and cryosectioned into 10 µm slices (Fig. 3a). Individual cells of the P*_gluD_::mNG* and P*_slpA_::mNG* reporter strains were readily seen (white arrows) in the colonic sections, with P*_slpA_::mNG* being the brighter reporter (Fig. 3b), whereas the P*_cwp2_::mNG* reporter strain was not reliably visualized (data not shown). Individual cells of all three *mScI3* reporter strains were also readily detectable, with the *mScI3* reporters providing a better signal-to-noise ratio than the *mNG* reporters due to lower levels of autofluorescence from plant fiber in the red channel.

**Fig. 3:**
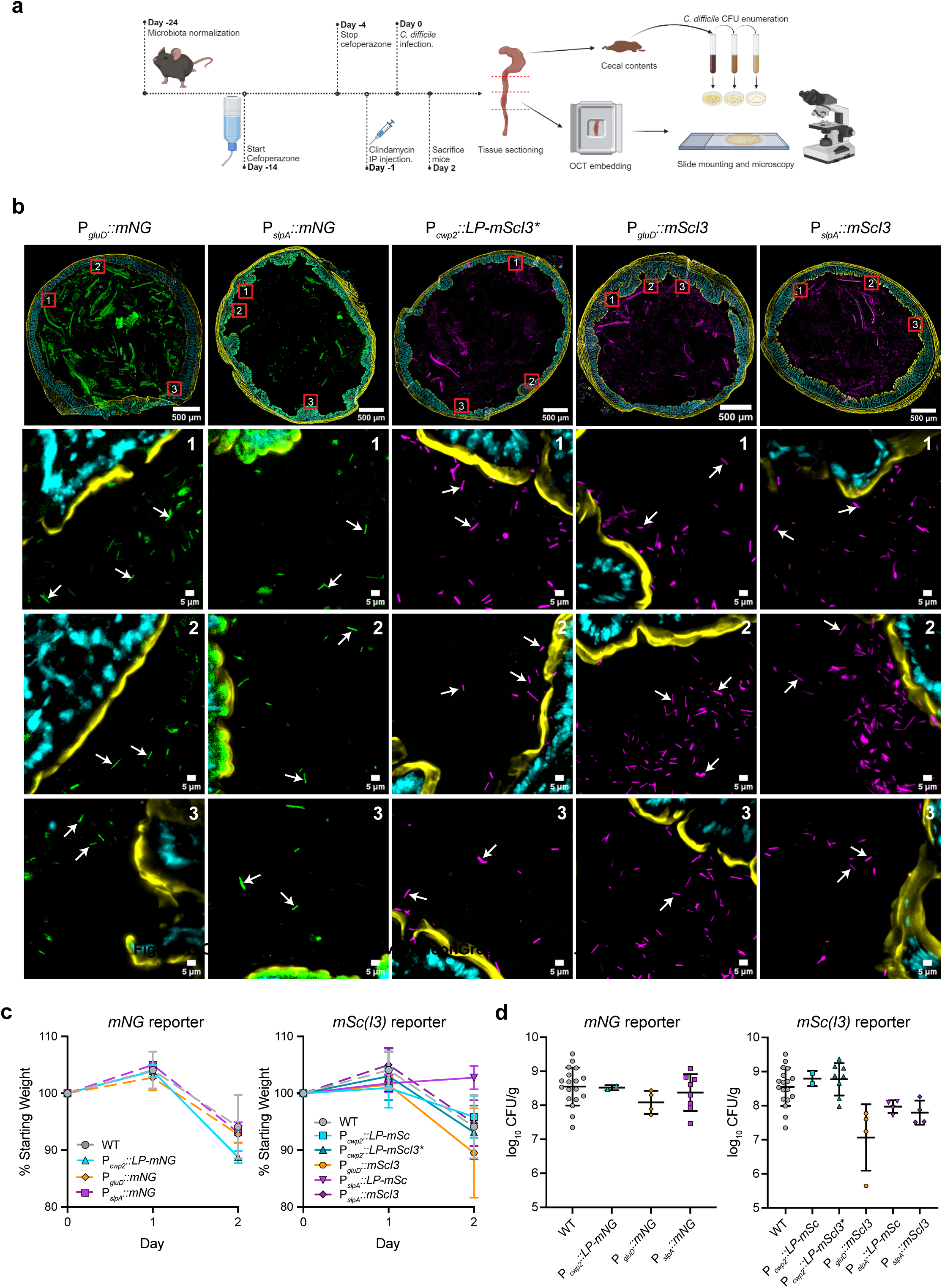
Validation of *mNeonGreen* and *mScarlet(I3*) reporters *in vivo*. **a.** Schematic of mouse infection timeline and downstream analyses including *C. difficile* CFU enumeration, mouse colonic tissue preparation, sectioning, and slide mounting for microscopy. Created with BioRender. **b.** Representative colonic section images of mice infected with constitutive reporter strains. Mice were infected with 1 x 10^6^ spores and sacrificed on Day 2 (48 hours) post-infection. Colon tissues were fixed in 4% PFA for 4 hours prior to cross-sectioning and embedding in OCT blocks. 10 µm cryosections were taken and stained with Phalloidin (F-actin, yellow) and DAPI (nuclei, cyan). Numbers in the top right of each inset panel indicate the corresponding image on the tissue cross-section. White arrows highlight *C. difficile* cells expressing either *mNG* (green) or *mS*(*I3*) (magenta). The P*_slpA_*::LP-*mSc* strain was not included in Fig. 3c because it failed to cause disease, and the P*_cwp2_::mNG* strains was also excluded because its fluorescence was too low to be reliably detected. **c.** Percent weight change of mice over the course of infection with the indicated *mNG* and *mSc*(*I3*) strains relative to the baseline weight measured at the time of infection (unmarked WT n = 23; P*_cwp2_::LP-mNG* n = 2; P*_gluD_::mNG* n = 4; P*_slpA_::mNG* n = 8; P*_cwp2_::LP-mSc* n = 2; P*_cwp2_::LP-mScI3\** n = 8; P*_gluD_::mScI3* n = 4; P*_slpA_::LP-mSc* n = 4; P*_slpA_::mScI3* n = 4). **d.** Cecal colony-forming units were measured on Day 2 (48 hours) post-infection. Lines show the geometric mean and standard deviation. (unmarked WT n = 18; P*_cwp2_::LP-mNG* n = 2; P*_gluD_::mNG* n = 4; P*_slpA_::mNG* n = 8; P*_cwp2_::LP-mSc* n = 2; P*_cwp2_::LP-mScI3\** n = 8; P*_gluD_::mScI3* n = 4; P*_slpA_::LP-mSc* n = 4; P*_slpA_::mScI3* n = 4).

Similar to a prior report imaging *C. difficile* during murine infection using FISH^36^, we found that most *C. difficile* cells were luminal, although cells in close association with the epithelium were readily detected, in contrast with this prior report^36^. *C. difficile* cells were also frequently found within a distinct border >50 µm above the epithelial layer (identified based on phalloidin staining); this localization was defined as mucosal (Fig. 3b). However, we were unable to directly visualize the mucus layer in these sections because preserving the mucus layer requires methacarn fixation, but this fixation method interferes with the fluorescence of the reporters^37,38^. Importantly, we observed these spatial distributions in multiple mice and multiple samples per mouse.

To ensure that the reporter strains were still capable of causing disease, we monitored the percent weight change of mice infected with *mNG* and *mSc*(*I*3) reporter strains relative to WT over the 48-hour infection period, as well as their colonization levels in the cecum when tissues were harvested 48 hours post-infection. Notably, the *mNG* strains caused similar levels of weight loss in mice (Fig. 3c) and colonized to similar levels as WT (Fig. 3d), indicating that the reporter constructs did not impair the infection process. Similar results were observed with *mSc*(*I3*) reporter strains, except that the P*_slpA_::LP-mSc* reporter strain failed to cause disease (Fig. 3c) and P*_gluD_::mScI3* and P*_slpA_::LP-mSc(I3)* strains colonized at ∼10-fold lower levels than WT (Fig. 3d).

### Identification of *mNeonGreen* and *mScarletI3* reporter strains with WT fitness *in vivo*

We next measured the fitness of *mScI3* reporter strains capable of causing disease relative to WT by infecting mice with 50:50 mixes of WT and the P*_cwp2_::LP-mScI3\**, P*_gluD_::mScI3*, or P*_slpA_::mScI3* reporter strains. Competitive indices were determined based on the ratio of red fluorescent cecal CFUs to WT non-fluorescent cecal CFUs. The P*_cwp2_::LP-mScI3\** and P*_gluD_::mScI3* strains exhibited equal fitness relative to WT, whereas the P*_slpA_::mScI3* strain had a slight fitness defect (∼3-fold, Fig. 4a). Similar levels of weight loss and colonization were observed for the competition infections as for the WT single-strain infection (Fig. 4b,c), except that the P*_gluD_::mScI3* vs. WT and P*_slpA_::mScI3* vs. WT competitions colonized to ∼5-10-fold lower levels.

**Fig. 4:**
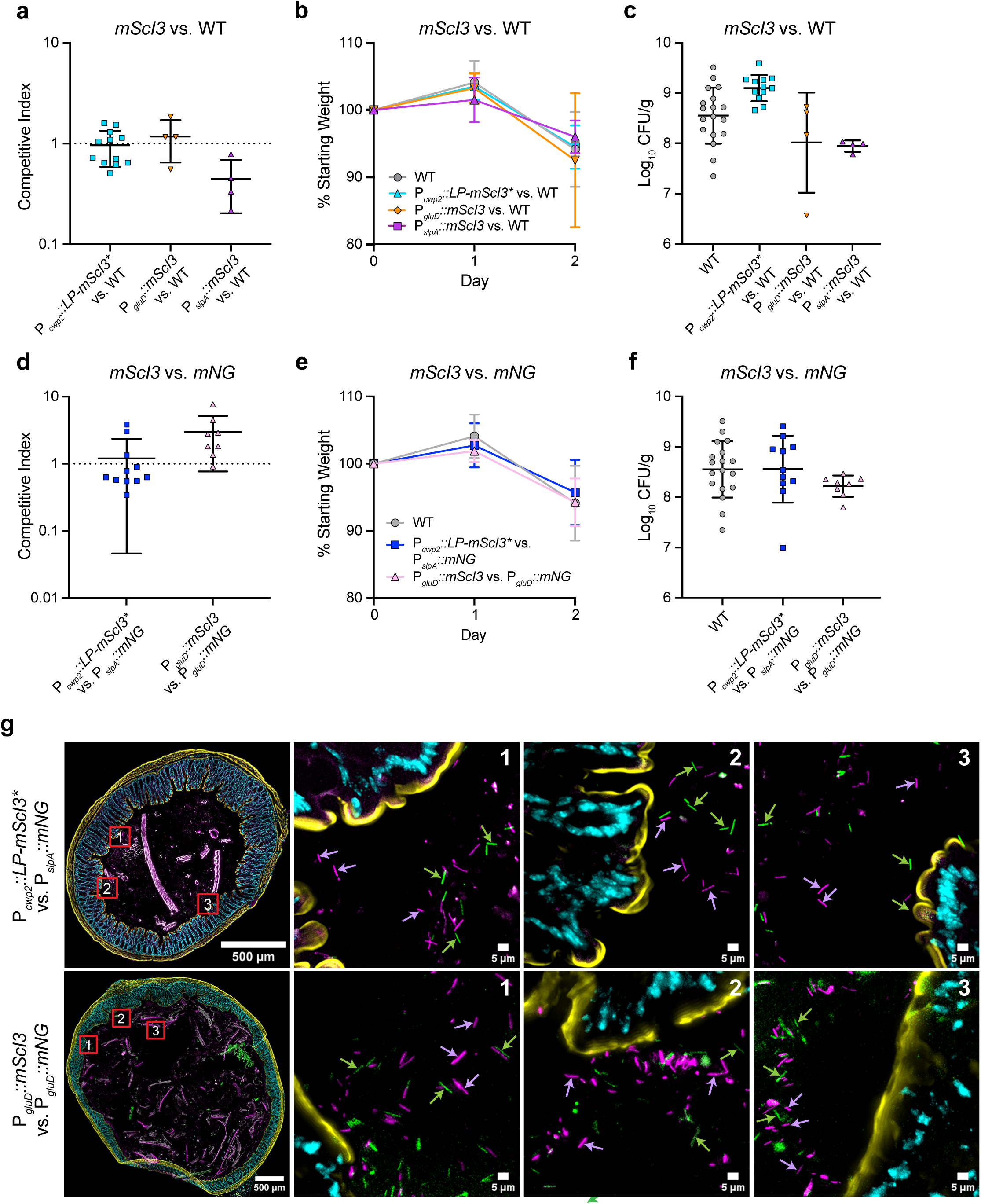
Relative fitness of *mNeonGreen* and *mScarletI3* reporter strains *in vivo*. **a, d.** Competitive indices of *mScI3*:WT or *mScI3*:*mNG* strains infections based on the red fluorescence of the resulting colonies. Lines show the geometric mean and standard deviation. (P*_cwp2_::LP-mScI3\** n = 12; P*_gluD_::mScI3* n = 4; P*_slpA_::mScI3* n = 4; P*_cwp2_::LP-mScI3\** vs. P*_slpA_::mNG* n = 11; P*_gluD_::mScI3* vs. P*_gluD_::mNG* n = 8; P*_cwp2_::LP-mScI3\** vs. P*_slpA_::mNG* n=11; P*_gluD_::mScI3* vs. P*_gluD_::mNG* n = 8). **b, e.** Percent weight change of mice over the course of infection with the indicated competition infections and a WT control. Competition infections used 5 x 10^5^ spores of each strain for a total infectious dose of 1 x 10^6^ spores per mouse; 1 x 10^6^ WT spores were used to infect mice as a control. Values are relative to the baseline weight measured at the time of infection (WT n = 23; P*_cwp2_::LP-mScI3\** vs. WT n = 12; P*_gluD_::mScI3* vs. WT n = 4; P*_slpA_::mScI3* vs. WT n = 4). **c,f.** Cecal colony-forming units measured Day 2 (48 hours) post-infection from the infections shown in **b** and **e**. Lines show the geometric mean and standard deviation. **g**. Representative images of mice co-infected with either P*_cwp2_::LP-mScI3\** and P*_slpA_::mNG* (top) or P*_gluD_::mScI3* and P*_gluD_::mNG* (bottom) from samples harvested Day 2 post-infection. 10 µm cryosections were stained with Phalloidin (F-actin, yellow) and DAPI (nuclei, cyan). Numbers in the top right of each inset panel indicate the corresponding image on the tissue cross-section. Green and mauve arrows indicate *C. difficile* cells expressing either *mNG* (green) or *mScI3* (magenta), respectively.

Since the auto-fluorescent nature of *C. difficile* in the green channel made it difficult to detect mNG-positive CFUs in a plate reader-based assay, we were unable to directly compare the fitness of *mNG* reporter strains to WT. Instead, we determined the relative fitness of P*_slpA_::mNG* and P*_gluD_::mNG* to P*_cwp2_::LP-mScI3\** and P*_gluD_::mScI3* reporter strains, respectively (Fig. 4d). Similar levels of weight loss and colonization were observed for the competition infections as with the WT infections alone (Fig. 4e,f). Similar fitness levels were observed for P*_cwp2_::LP-mScI3\** and P*_slpA_::mNG* strains, whereas the P*_gluD_::mScI3* strain had a ∼4-fold fitness advantage over P*_gluD_::mNG* (Fig. 4d).

Importantly, *mScI3* and *mNG* reporter cells were readily detected in colonic sections, and they exhibited equivalent spatial distributions in the lumen and close to the colonic epithelium, with green and magenta cells being similarly intermixed in these microenvironments (Fig. 4g). However, it was harder to detect the P*_gluD_::mNG* strain because its signal is weaker relative to P*_slpA_::mNG* (Fig. 4g). Taken together, these analyses identified P*_cwp2_::LP-mScI3\** and P*_slpA_::mNG* reporters as the best red and green constitutive reporter strains to visualize during infection.

### Development of dual reporter strains for visualizing phenotypic heterogeneity in toxin gene expression *in situ* during *C. difficile* infection

We next combined our optimized constitutive reporters with a toxin gene reporter to visualize the subset of cells expressing toxin genes *in situ* during infection. Specifically, we generated a dual reporter strain carrying the constitutive P*_cwp2_::LP-mScI3* reporter with a toxin gene-specific *mNG* reporter that we previously constructed, P*_tcdA_::mNG*^32^. The reporters were integrated into the chromosomes of WT, Δ*tcdR*, and Δ*rstA* strains at a neutral locus to assess the ability of the reporter system to read out toxin gene expression at the single-cell level. TcdR is the toxin gene-specific sigma factor that activates toxin gene expression, so toxin gene expression is essentially null in a *ΔtcdR* strain *in vitro*^32,39^. Conversely, RstA is a negative regulator of toxin gene expression, so a Δ*rstA* strain over-expresses toxin genes^40^ and exhibits a higher frequency of toxin gene expression *in vitro*^32^.

We first validated the bimodal nature of toxin gene expression in the dual reporter strains after overnight growth in TY broth (Extended Data Fig. 1a; dual reporter strains are designated with an * from herein). While the constitutive *mScI3* reporter exhibited uniform fluorescence regardless of strain background (Extended Data Fig. 1b), *tcdA* toxin gene expression was bimodal, being observed in ∼37% of cells in the WT* background (Extended Data Fig. 1c, d). Toxin gene expression was ∼2-fold higher (74%) and more frequent in the *ΔrstA\** strain than in WT* (Extended Data Fig. 1c, d), consistent with our prior analyses of toxin gene expression with single reporter strains^32^.

When the dual reporter strains were used to infect mice, WT* and Δ*tcdR\** colonized to similar levels, whereas Δ*rstA\** colonized at ∼10-fold lower levels than WT* on Day 2 post-infection (Fig. 5a). Despite this reduced colonization level on Day 2, the Δ*rstA\** reporter strain nevertheless caused more weight loss in mice than the WT* reporter strain and unmarked WT strain (Fig. 5b). This virulence phenotype is consistent with prior work showing that an *rstA* mutant strain causes greater disease in hamsters, likely due to its higher toxin production, and colonizes to lower levels^41^. As expected, the Δ*tcdR* strain failed to cause weight loss (Fig. 5b).

**Fig. 5:**
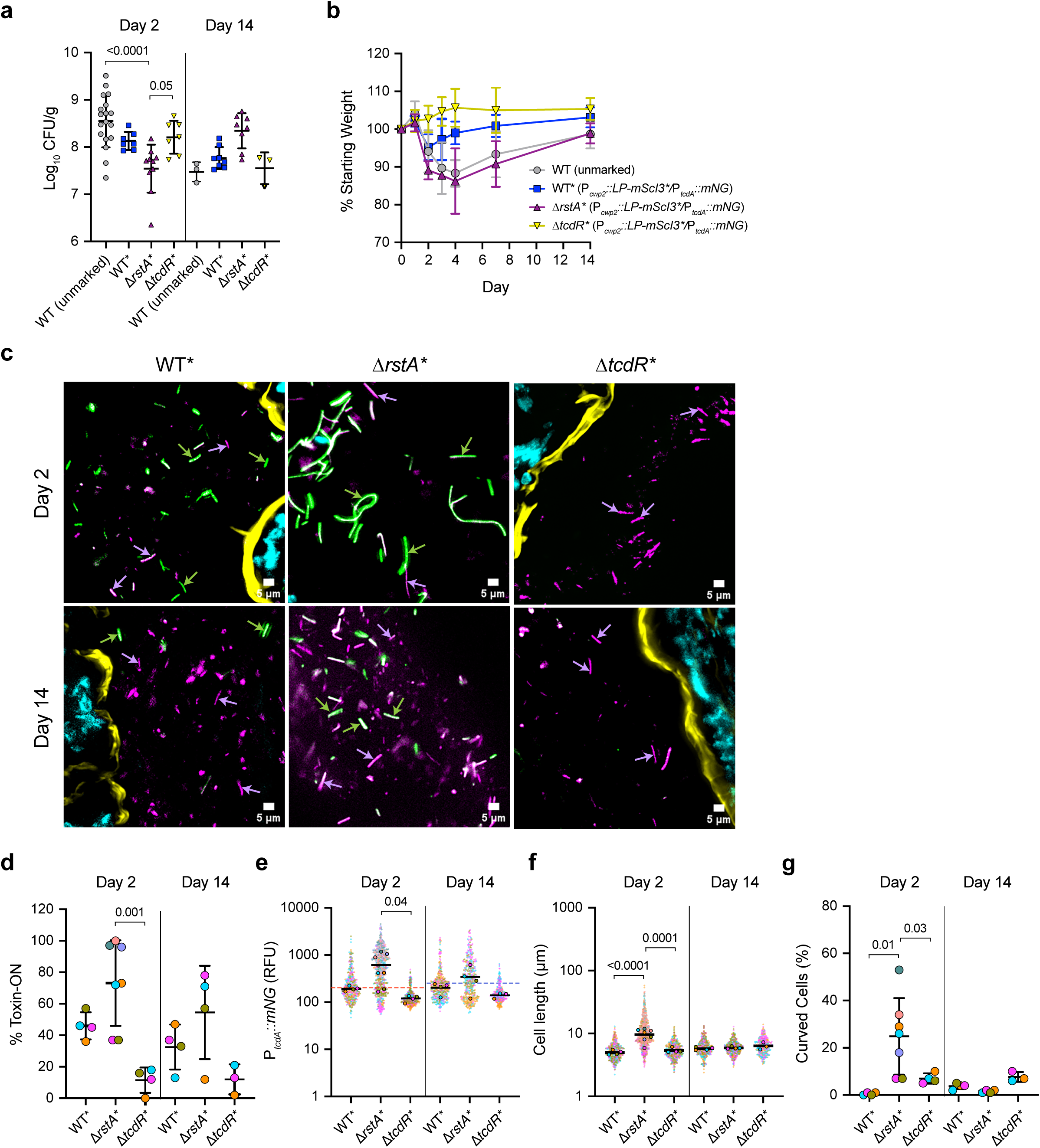
*In situ* analyses of phenotypic heterogeneity in toxin gene expression during murine infection reveals that a toxin gene-overexpressing mutant forms filaments during the acute phase of infection. **a.** Cecal colony-forming units measured either Day 2 (WT n = 18; WT*n = 7; Δ*rstA*\* n = 10; Δ*tcdR\** n = 7) or Day 14 (WT n = 3; WT* n = 8; Δ*rstA*\* n = 8; Δ*tcdR\** n = 73, * = P*_cwp2_::LP-mScI3*/P*_tcdA_::mNG*) post-infection. Lines show the geometric mean and standard deviation. **b.** Percent weight change of mice over the course of infection with the indicated dual reporter strains. Mice were infected with 1 x 10^6^ spores of the indicated *C. difficile* strains. Values are relative to the baseline weight measured at the time of infection (WT n = 23; WT* n = 8; Δ*rstA*\* n = 11; Δ*tcdR*\* n = 4). **c.** Representative images of colonic sections harvested from mice at the indicated time for WT*, Δ*rstA\**, or Δ*tcdR\** dual reporter strains. Colon tissues were fixed in 4% PFA for 4 hours prior to cross-sectioning and embedding in OCT blocks. 10 µm sections were stained with Phalloidin (F-actin, yellow) and DAPI (nuclei, cyan). Green arrows indicate the subset of Toxin-ON *C. difficile* cells; mauve arrows indicate Toxin-OFF cells, where only the constitutive *mScI3* reporter was detectable (magenta). **d.** Percentage of cells expressing the toxin-specific P*_tcdA_::mNG* reporter in the indicated dual reporter strains in colonic sections from mice. The % Toxin-ON was determined by the proportion of cells with mNeonGreen signal greater than one standard deviation above the mean fluorescence of the Δ*tcdR\** dual reporter strain (indicated by dashed red line Day 2 post-infection; blue line Day 14 post-infection). The mean and standard deviation are shown based on the percentages measured for four mice, except for Δ*rstA\**, where seven mice were analyzed. One hundred cells were quantified per mouse. **e.** Superplot of toxin-specific reporter fluorescence (P*_tcdA_::mNG*) at the single-cell level in colonic sections of mice infected with the indicated dual reporter strains Day 2 or Day 14 post-infection. **f.** Superplot of cell lengths measured for individual cells detected in colonic sections for WT*, Δ*rstA\**, and Δ*tcdR\** dual reporter strains. For all superplots, the median value of 100 cells per mouse was determined (colored outlined dot); the median values were then averaged to give the mean relative fluorescence unit (RFU) or cell length (horizontal black line) (n = 400 cells per condition, Δ*rstA\** at Day 2, n = 700). **g.** Percentage of curved cells for WT*, Δ*rstA\**, and Δ*tcdR\** dual reporter strains. The mean and standard deviation are shown based on the percentages measured for a minimum of four mice (100 cells were quantified per mouse). Statistical significance was determined using a one-way ANOVA and Tukey’s test. Only statistically significant comparisons are shown.

To determine whether the reduced cecal CFU levels observed with the *ΔrstA\** reporter strain were specific to acute infection or whether this strain exhibits colonization defects at later time points, we analyzed the infection dynamics of the dual reporter strains over the course of a 14-day infection. While the Δ*rstA\** reporter strain and unmarked WT strain caused similar levels of weight loss (10–20% of initial body weight) during peak disease severity between days 2 and 4, the WT* reporter strain induced less weight loss (∼5%) (Fig. 5b). Notably, by Day 14, the Δ*rstA\** reporter strain achieved ∼3-fold higher colonization levels than both WT strains tested (p < 0.01), despite colonizing at lower levels on Day 2 (Fig. 5a, p < 0.0001).

Since the WT* reporter strain caused less weight loss than the unmarked WT strain on Days 3 and 4 post-infection, we sought to assess whether the Δ*rstA\** reporter strain behaves differently from its unmarked parental Δ*rstA* strain by analyzing the weight loss of unmarked Δ*rstA* and an Δ*rstA*/*rstA* complementation strain. Importantly, the Δ*rstA* dual reporter strain caused similar levels of weight loss as the unmarked strains, indicating that the dual reporter does not affect the virulence of the Δ*rstA* strain. (Extended Data Fig. 2).

When toxin gene expression in the dual reporter strains was analyzed in colonic sections taken from tissues harvested on Day 2 and Day 14 post-infection, toxin gene expression (i.e., production of mNG per cell) in both the WT* and Δ*rstA\** strains was found to be heterogeneous during infection (Fig. 5c,d), with a higher proportion of Δ*rstA\** reporter strain cells expressing toxin genes than WT* at Day 2 (73% vs. 46%, respectively, Fig. 5d), similar to our *in vitro* analyses (Extended Data Fig. 1). As expected, the constitutive *mScI3\** reporter signal was consistent across the dual reporter strains and across days 2 and 14 (Extended Data Fig. 3). When we compared the magnitude of toxin gene expression in the subset of cells identified as Toxin-ON, the Δ*rstA\** reporter strain expressed toxin genes at ∼2-3-fold higher levels on average than the WT* reporter strain on Day 2 (Fig. 5e). Interestingly, the average magnitude of toxin gene expression for Toxin-ON cells was lower on Day 14 for the Δ*rstA\** strain relative to Day 2. While Day 2 and Day 14 samples were harvested from different mice due to the endpoint nature of the assay, this trend was observed in two independent experimental infections (Fig. 5e). In contrast, there was little change in the magnitude of toxin gene expression for WT* cells between Day 2 and Day 14, although there was a slight decrease in the proportion of WT* cells expressing toxin genes on Day 14 (Fig. 5d, e). Surprisingly, about 10% of Δ*tcdR\** cells were identified as Toxin-ON in these analyses based on the expression of toxin genes one standard deviation above the mean determined based on single-cell measurements across a minimum of four mice. While the magnitude of toxin gene expression in Toxin-ON Δ*tcdR\** cells was low compared to the WT* and Δ*rstA\** strains, mechanisms for inducing toxin gene expression independent of TcdR appear to exist *in vivo* (Fig. 5d,e) since this Toxin-ON population is not observed during *in vitro* growth (Extended Data Fig. 1a,c,d).

### Relationship between toxin gene expression and spatial localization during infection

Since toxin production changes the metabolic environment surrounding *C. difficile* by liberating nutrients close to the epithelium^42^, we asked whether toxin production impacts the localization of *C. difficile* within the colon. We found that the Δ*tcdR\** strain exhibited similar spatial distributions in colonic sections relative to WT* and Δ*rstA\** strains, with Δ*tcdR* being readily observed close to the epithelium (Fig. 5c, Extended Data Fig. 4). Thus, higher toxin gene expression does not appear to affect the proximity of *C. difficile* cells to the epithelium, at least under the conditions analyzed.

Since toxin gene expression is responsive to many environmental inputs^39^, we assessed whether the magnitude and proportion of toxin gene-expressing cells are spatially or temporally regulated during infection. To this end, we grouped cells into three spatial categories, the lumen, mucosa, or epithelium, and measured the fluorescence of the P*_tcdA_::mNG* reporter and quantified the magnitude and frequency of toxin gene expression (Extended Data Fig. 5,6). These analyses revealed that neither the proportion of toxin gene-expressing cells nor the magnitude of their gene expression is affected by the proximity of *C. difficile* to the epithelium (Extended Data Fig. 5,6).

### Discovery of a filamentous cell morphology during acute infection by a mutant that over-expresses toxin genes

While no differences in the spatial distribution of *C. difficile* were observed between the WT*, Δ*rstA\**, and Δ*tcdR\** strains, we nevertheless discovered that the Δ*rstA* strain frequently forms curvy, filamentous cells on Day 2 (Fig. 5c,f,g Extended Data Fig. 4). Indeed, the Δ*rstA\** reporter strain was 2-fold longer on average than the WT* or Δ*tcdR\** reporter strains, with some cells exceeding over 50 µm. Notably, Δ*rstA\** cells were more likely to be curved (25%) compared to WT* (0.5%) and Δ*tcdR\** (7%) on Day 2, when peak weight loss is often observed (Fig. 5f,g). However, by Day 14, the percentage of curved Δ*rstA\** cells decreased to 1.5% similar to WT (Fig. 5g).

Since the abnormal cell morphologies observed for the *ΔrstA\** reporter strains could be due to the over-expression of *mNG* in Δ*rstA\** cells during peak infection, we analyzed the morphology of a Δ*rstA* strain expressing only the constitutive *mScI3* reporter (Extended Data Fig. 7). The *ΔrstA* single constitutive reporter strain was shorter than the *ΔrstA\** dual reporter strain, but it was ∼20% longer on average than the WT single reporter strain, although this difference was not statistically significant (Fig. 5f, Extended Data Fig. 7a). Notably, the *ΔrstA* single reporter strain showed a trend toward forming more curved cells relative to the WT single reporter strain, similar to analyses of the Δ*rstA* dual reporter strain to WT and Δ*tcdR* dual reporter strains (Extended Data Fig. 7b). These data imply that the curvy, elongated cell morphology is unique to the Δ*rstA* strain during acute stages of infection. This altered morphology appears to be linked to high-level toxin gene expression because we found that cells expressing P*_tcdA_::mNG* to high levels were more likely to be elongated in the *ΔrstA* strain (Extended Data Fig. 8). Collectively, these data indicate that high level mNG production exacerbates the morphological changes in the Δ*rstA* reporter strain, but it is not the sole driver of this phenotype.

Since the elongated morphology of the Δ*rstA\** strain was only observed during the acute stage of infection, we sought to determine whether different phases of growth *in vitro* or minimal media conditions could also induce this phenotype. To this end, we grew WT, Δ*rstA*, and Δ*tcdR* dual reporter strains in nutrient-rich TY or minimal CDDM media and imaged cells at mid-log or stationary phase. No morphological differences across strains or conditions were detected (Extended Data Fig. 9a,b), and the frequency of toxin gene expression in each strain was similar between log and stationary phase, as well as between the different media (Extended Data Fig. 9d). Thus, the abnormal morphology of the Δ*rstA*\* strain appears to be specific to the acute phase of murine infection.

## Discussion

Phenotypic heterogeneity allows bacteria to maximize their fitness in dynamic environments^43^ such as those found in the gut^15,44^, but it has been challenging to image this heterogeneity at the single-cell level in the dense colonic environment *in situ*. Here, we developed chromosomally-encoded, spectrally-compatible fluorescent reporters that allow us to visualize the subset of *C. difficile* cells expressing toxin genes *in situ* during murine infection. While this is the first report of phenotypic heterogeneity being visualized *in situ* in the colon during a natural infection to our knowledge, our approach could be applied to study heterogeneity in gene expression *in situ* for the many colonization and virulence factors that *C. difficile* expresses in a bimodal manner^25,32,33,45^.

Our analyses provide novel insight into the spatial and temporal dynamics of toxin gene expression during *C. difficile* infection. First, they revealed that, while *C. difficile* is primarily a luminal pathogen, a sub-population is proximal to the epithelial layer, and a minority of this sub-population directly interacts with the epithelium (Figs. 3-5). Second, a cell’s location relative to the colonic epithelium did not affect the magnitude or frequency of toxin gene expression, and toxin gene expression did not grossly affect where *C. difficile* localized within the colon (Extended Data Fig. 5,6). Third, toxin gene expression frequency and magnitude did not change over time except for in the Δ*rstA* strain, where toxin genes were expressed at lower levels and less frequently in the *ΔrstA* strain 14 days post-infection compared to 2 days post-infection (Fig. 5d,e). This could represent an adaptation by the Δ*rstA* strain or a selection for cells expressing toxin genes at lower levels.

Based on these observations, *C. difficile* likely uses heterogeneity in toxin gene expression as a bet-hedging strategy to partition the metabolically intensive task of toxin production so that non-toxin-producing cells, and ultimately the entire population, can benefit^25,44,46^. Specifically, the ability of toxin-producing cells to damage the host epithelium and release nutrients likely benefits non-toxin-producing cells by freeing this sub-population from the metabolic costs of toxin synthesis. Non-toxin-producing cells may then allocate resources toward processes such as specific metabolic programs or sporulation, a critical step in disease dissemination. Indeed, we previously reported a division of labor between toxin production and sporulation during the growth of *C. difficile* under certain conditions *in vitro*^32^. Consistent with this hypothesis, the Δ*rstA* mutant’s reduced colonization levels (Fig. 5a) could suggest that over-producing toxin decreases *C. difficile*’s ability to expand in the gut during the acute phase. Alternatively, it may also reflect filamentous cells producing fewer colonies on a plate. Regardless, given RstA’s multiple roles in regulating sporulation, toxin production, and motility^40,41^, testing our hypothesis would require mutants that specifically over-express toxin genes.

Our analyses also highlight a key benefit of localizing bacteria at the single-cell level *in situ* because we serendipitously discovered that the Δ*rstA* strain adopts a filamentous, curved morphology specifically during the acute phase of infection. This abnormal morphology may be due to stressful conditions encountered during this infection phase because Δ*rstA* cells grown in TY or minimal media broth exhibit WT cell morphology despite expressing toxin genes (and the *mNG* reporter gene) to high levels (Extended Data Fig. 9). The specific conditions responsible for this phenotype remain unclear, but stresses that are present during infection, such as oxidative stress, induce DNA replication arrest through *C. difficile’s* SOS response, which can cause cell filamentation^47^.

Notably, our identification of spectrally compatible constitutive reporter strains with equivalent fitness (P*_cwp2_::LP-mScI3\** and P*_slpA_::mNG*) will enable future studies focused on identifying genes that impact the relative spatial distribution of *C. difficile* cells in the colon epithelium. Indeed, our analyses are the first to show that a subpopulation of *C. difficile* is found in close proximity to the epithelium (Figs. 3-5). This previously unappreciated subpopulation could be important for delivering *C. difficile*’s toxins more efficiently to their receptors or establishing biofilms for promoting recurrent infections. Since genes important for this newly appreciated localization to the epithelium remain unknown, our reporter system should facilitate the identification of these genes by analyzing the relative distribution of *mScI3* and *mNG* reporter strains in different mutant backgrounds during single or competitive infections. The constitutive reporter pairs can also be used to analyze founder effects^17,48^, such as whether avirulent strains can outcompete virulent strains by inhabiting different spatial niches at early stages of infection^49^ or whether two strains exhibit different distributions during long-term colonization or recurrent infections.

Finally, since promoters in other clostridia are efficiently used in *C. difficile* and vice versa^50–54^, our approach opens the door for investigating not only the spatial and temporal regulation of gene expression during infection by *C. difficile* but also other clostridial organisms that modulate gut health or disease. Given the critical role of clostridial organisms in maintaining gut health^55^, the ability to analyze the spatial and temporal dynamics of their colonization and gene expression will enable mechanistic insights into their impacts on the gut microbiome and human health.

## Materials and Methods

### Bacterial strains and growth conditions

All *C. difficile* strains used for this study are listed in Table S1. All constructed strains derive from the sequenced clinical isolate 630, with the erythromycin-sensitive 630Δ*erm*Δ*pyrE* serving as the parental strain for all strain constructions using *pyrE*-based allele-coupled exchange (ACE)^57^. Strains were grown on brain heart infusion supplemented with yeast extract and cysteine (BHIS); taurocholate (TA; 0.1% [wt/vol]; 1.9 mM), cefoxitin (8 mg/mL), thiamphenicol (10 to 15 mg/mL), and kanamycin (50 mg/mL) were added as needed. For ACE, the *C. difficile* defined medium (CDDM)^58^ was supplemented with 5-fluoroorotic acid (5-FOA; 2 mg/mL) and uracil (5mg/mL). All *C. difficile* strains were first grown overnight from glycerol stocks on BHIS plates supplemented with TA (0.1% [wt/vol]).

*Escherichia coli* strains used for HB101/pRK24-based conjugations are listed in Table S1. *E. coli* strains were grown at 37°C with shaking at 225 rpm in Luria-Bertani (LB) broth. The medium was supplemented with ampicillin (50mg/mL) and chloramphenicol (20mg/mL) as needed.

### *C. difficile* strain construction

Single-reporter strains were generated by conjugating HB101 carrying pMTL-YN1C plasmids into Δ*pyrE*-based strains using ACE. To generate dual-reporter strains, constitutive reporters were introduced downstream of the *sipL* locus of 630Δ*erm*Δ*pyrE*, 630Δ*erm*Δ*tcdR*Δ*pyrE,* or 630Δ*erm*Δ*rstA*Δ*pyrE* using ACE and pMTL-YN3-P*_cwp2_*::*LP-mScarlet(I3)*. The second reporter, P*_tcdA_*::*mNeonGreen*, was then introduced into the *pyrE* locus of the resulting strains using pMTL-YN1C-P*_tcdA_*::*mNeonGreen*. At least two clones of each strain generated by allelic exchange were phenotypically characterized prior to restoration of the *pyrE* locus using pMTL-YN1C.

### Growth curves

*C. difficile* cultures were grown in 2 mL BHIS medium to stationary phase and then were back-diluted 1:50 in 2 mL BHIS and grown until an OD_600_ of 0.5 was reached. All strains were normalized to an OD_600_ of 0.5 if growth rates varied. Next, strains were diluted into 200 µl of BHIS broth in a 96-well plate to a starting OD_600_ of 0.05. Each strain was grown in technical triplicate alongside BHIS blanks. Plates were read in an Epoch plate reader (BioTek) in the anaerobic chamber with OD_600_ readings performed every 15 min after a 2-min linear shake. For *in vitro* toxin visualization, *C. difficile* cultures were grown in 2 mL TY broth to mid-log phase, then back-diluted 1:50 in 2 mL TY and grown overnight followed by fixation and microscopy.

### Cell fixation of broth-grown cultures

Cells were grown to mid-log phase in BHIS broth or to stationary phase in TY broth overnight. Next, 500 uL of culture was added directly to a tube containing 120 μL of a 5× fixation cocktail (100 μL of 16% [wt/vol] paraformaldehyde aqueous solution [methanol-free]) and 20 μL of 1 M NaPO_4_ buffer (pH 7.4). The samples were mixed and incubated aerobically for 30 min at room temperature in the dark followed by 30 min on ice in the dark. The fixed cells were washed 3 times in phosphate-buffered saline (PBS) and resuspended in 500 μL (depending on the density of the culture). Two μL of fixed cells were spotted onto a 1.5% agarose pad. Once the spot of cells was dry, a coverslip was added to seal the slide, and the slide was incubated at 37°C for 2 hours to allow for fluorophore maturation before imaging.

### Murine infections

Seven-week-old female C57BL/6 cage-mate mice (Jackson Laboratory) were housed together in a large sterile rat cage to allow microbiota normalization over a 10-day period. Mice were fed irradiated Lab Diet 2918 throughout. After 10 days, cefoperazone (0.5 mg/mL) was added to the drinking water, which was provided ad libitum for another 10 days. Mice were then returned to sterile drinking water for 2 days before receiving a single intraperitoneal injection of clindamycin (10 mg/kg in 200 µL), prepared in 1× PBS and filter-sterilized.

Twenty-four hours after clindamycin administration, mice were moved from the large rat cage into smaller mouse cages, housing 4 mice per cage, and were inoculated via oral gavage with 1×10⁶ *C. difficile* spores in 200 µL of 1× PBS. At the time of oral gavage, baseline weights were recorded by transferring mice using large forceps into sterile plastic Nalgene cups and weighing them on a digital scale inside a biosafety cabinet.

Following inoculation, mice were monitored daily and weighed using the same procedure for either 2 or 14 days. At the experimental endpoint, mice were euthanized by isoflurane anesthesia followed by cervical dislocation. Tissues were collected as needed.

### CFU enumeration

Upon sacrifice, cecal contents were collected into pre-weighed 1.5 mL Eppendorf tubes for *C. difficile* biomass quantification. The tubes were reweighed to calculate the mass of the collected content. Tubes were then transferred into an anaerobic chamber, where 1 mL of pre-reduced 1× PBS was added to each to resuspend the cecal content. After thorough homogenization, 10 µL of the suspension was added to 90 µL of 1× PBS in the first well of a 96-well plate to begin a series of seven 10-fold serial dilutions. For each subsequent dilution, 10 µL was transferred into 90 µL of PBS in the next well, mixing thoroughly at each step. From each dilution, 5 µL was spotted onto TCCFA plates to select for *C. difficile*. Plates were incubated for 24 hours in the anaerobic chamber, after which colony-forming units (CFUs) were counted and normalized to the mass of the original cecal content.

### Competition assays

Mice were infected with a total of 1×10⁶ *C. difficile* spores, consisting of a 1:1 mixture of two strains: either wild-type and a fluorescent reporter strain, or two distinct fluorescent reporter strains (mNeonGreen and mScarlet-I3). Infections were performed as described above, and mice were sacrificed 48 hours post-infection. Cecal contents were collected, weighed, and processed as previously described. Ten-fold serial dilutions were prepared, and 100 µL of dilutions 4–7 were plated onto TCCFA plates to select for *C. difficile*.

After 24 hours of anaerobic incubation, single colonies were picked using a pipette tip and resuspended in 200 µL of 1×PBS in 96-well plates (one plate per mouse). Plates were incubated at 37°C for at least one hour to allow fluorophore maturation. Fluorescence was then measured using a Synergy H1 Microplate Reader (excitation: 570 nm, emission: 600 nm). Wells with a positive fluorescence signal were identified as *mScarlet-I3*-expressing colonies, while wells lacking signal represented either wild-type or mNeonGreen-expressing colonies. The ratio of mScarlet-I3 to mNeonGreen colonies recovered from each mouse was normalized to the input spore ratio.

### Mouse colon tissue preparation

After mice were sacrified, the entire colon was removed and placed into a 4% wt/vol paraformaldehyde and 1xPBS solution for 4 hours. Following the 4-hour tissue fixation, cross-sections of tissue were prepared by cutting 5mm transverse rings. The tissues were then placed in O.C.T Compound (Tissue-Tek) in Seal’N Freeze Cryotray tissue cassettes (Electron Microscopy Sciences) and flash frozen in a dry ice 100% ethanol bath. Frozen blocks were then processed by the Tufts Animal Histology Core, where 10 μm sections were cryosectioned and mounted to slides. Any remaining tissue that was not embedded was placed in 1xPBS for long-term storage. Mounted slides were kept at −80°C until they were ready to be stained and imaged.

### Staining mouse colon slides

Mounted slides were removed from the freezer and placed on the bench at room temperature for 30 minutes, and then an ImmEdge Pen (Vector Laboratories) hydrophobic marker was used to trace around the tissue sections to facilitate staining of the sections. Slides were blocked with blocking buffer (1xPBS, 3% BSA [w/v], 0.1% Triton X-100) for 30 minutes. Blocking buffer was then discarded, and stain containing blocking buffer, DAPI (100 ng/mL), and Phalloidin CruzFluor™ 647 (1:100) were added for 45 minutes. Following staining, slides were washed with blocking buffer 3X for 5 minutes before mounting with Prolong Diamond mounting medium. Slides were left to cure overnight at room temperature before imaging the following day.

### Fluorescence microscopy

All images were acquired using a Leica DMi8 Thunder imager equipped with an HC PL APO 63×/1.4 numerical aperture (NA) phase-contrast oil immersion objective (*in vitro* microscopy) or an HC PL APO 20x/0.80 dry objective (mouse tissue sections). Excitation light was generated by a Lumencor Spectra-X multi-LED light source with integrated excitation filters. For all fluorescent channels, an XLED-QP quadruple-band dichroic beam-splitter (Leica) was used along with an external filter wheel (Leica). Phase-contrast images were taken with a 50-ms exposure time. mScarlet(I3) was excited at 550 nm (33% intensity) with 200 ms exposure time, a dichroic mirror at 570nm, and emitted light was filtered using a 595/40 nm emission filter (Leica). mNeonGreen was excited at 470 nm with a 300 ms exposure, a dichroic mirror at 419 nm, and emitted light was filtered using a 515/40 nm emission filter. Emitted and transmitted light was detected using a Leica DFC 9000 GTC sCMOS camera. All strains for a given experiment were spotted and captured sequentially on the same agarose pad.

For imaging mouse colonic tissue, stained slides were imaged using the same Leica microscope as indicated above using a 20x dry objective lens. DAPI stain for visualizing colonic epithelial cell nuclei was excited at 395 nm (10% intensity) with a 50-ms exposure, dichroic mirror at 415 nm, and emitted light was filtered using a 430/36-nm emission filter (Leica).. The phalloidin F-actin stain was excited at 640 nm, a dichroic mirror at 660 nm, and emitted light was collected at 720/50 nm. Cells containing either mScarlet(I3) or mNeonGreen fluorescent proteins were imaged as described above. 5 µm step sizes were used to images through the colon tissue.

### Image analysis and quantification

After image acquisition, images were processed using Leica Instant Computational Clearing (ICC) to eliminate bleed-through of fluorescent signals into neighboring cells. The adaptive strategy was executed with a feature scale of 2,500 nm and 98% strength. Following ICC, images were exported from Leica’s LASX software and further processed in FIJI. Specifically, images were cropped to remove out-of-focus regions, and the best-focused Z-plane was selected for each channel to correct for chromatic aberration.

Quantitative analysis of broth-grown cells was performed using the SuperSegger^56^ pipeline in MATLAB with the provided 60× analysis settings. The resulting clist matrices, which contained single-cell data, were exported from MATLAB. For display purposes, image scaling was uniformly adjusted for brightness and contrast across all strains in each experiment (unless marked “ADJUSTED” on the image). At least three images per strain were captured in each replicate, and every strain was analyzed using three biological replicates, with image analysis performed on at least two positions per replicate.

To quantify the proportion and magnitude of *C. difficile* toxin gene expression *in situ* during infection, *C. difficile* cells were first identified using the mScarlet-I3 channel. Median fluorescence intensity for each cell was then measured in both the mScarlet-I3 and mNeonGreen channels. This approach minimizes selection bias by including all identified cells, not just those with the highest mNeonGreen signal. Individual cells were manually traced along their full length using segmented lines in FIJI. For each experimental group, cross-sectional images of the colon were analyzed from at least three different mice. A minimum of 100 cells per image was measured.

To determine whether a cell was classified as “Toxin-ON,” the mNeonGreen signal from a toxin-null Δ*tcdR* dual reporter strain was used as a baseline. Cells with fluorescence values greater than one standard deviation above the mean of this baseline were defined as Toxin-ON. Thresholds were determined separately for Day 2 and Day 14 post-infection.

For spatial localization analysis, cells were categorized into three groups: luminal, mucosal, and epithelial. Luminal cells were defined as those located in the central lumen of the colonic cross-section. Mucosal cells formed a distinct border adjacent to the epithelium (> 50 µm), as determined by F-actin phalloidin staining. Cells that were in direct contact with the epithelial layer, i.e., where the mScarlet-I3 signal co-localizes with phalloidin, were classified as epithelial-associated.

### Cell curvature and length measurements

To assess cell curvature, segmented lines were manually drawn along the length of individual *C. difficile* cells using FIJI (ImageJ) based on the mScarlet-I3 fluorescence signal from reporter strains. The total length of each segmented line was recorded to represent the cell’s actual path length. X and Y coordinates of the cell endpoints were extracted and used to calculate the Euclidean distance between the two ends of each cell using a FIJI plugin. A curvature ratio was then calculated by dividing the total segmented length by the Euclidean distance. Cells with curvature ratios greater than 1.03 were classified as curved.

---

**Extended Data Figure 1. Toxin gene expression in dual reporter strains during growth in broth culture.**

**Extended Data Figure 2. Percent weight change of mice over the course of infection with the indicated dual reporter or unmarked strains.**

**Extended Data Figure 3. Quantification of constitutive reporter fluorescence during murine infection.**

**Extended Data Figure 4. Representative images of colonic sections harvested from mice for dual reporter strains.**

**Extended Data Figure 5. Spatial distribution of toxin gene expression during murine infection.**

**Extended Data Figure 6. Spatial distribution of toxin gene expression in individual mice.**

**Extended Data Figure 7. Comparison of the cell morphology of dual vs. single reporter strains during infection.**

**Extended Data Figure 8. Relationship between toxin gene expression and cell length during murine infection.**

**Extended Data Figure 9. Toxin gene expression in dual reporter strains during growth in nutrient-rich TY or minimal CDDM broth at mid-log or stationary phase.**

