## Supplemental Figures for "Visualizing phenotypic heterogeneity and single-cell morphology *in situ* during gut infection"

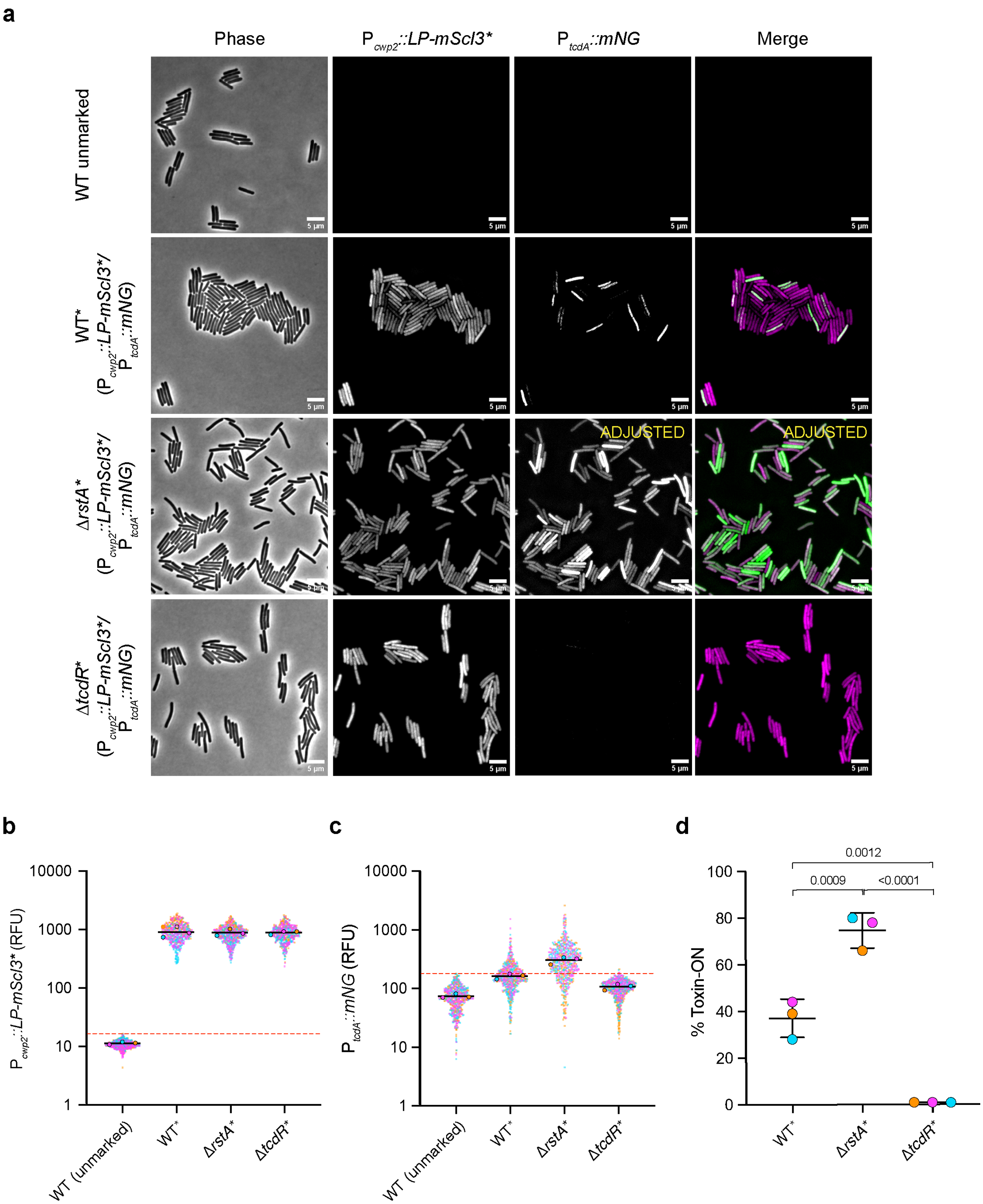


**Extended Data Figure 1. Toxin gene expression in dual reporter strains during growth in broth culture. a.** Representative fluorescence microscopy of unmarked WT or dual reporter strain cells (* = $P_{cwp2}$*::LP-mScI3*/$P_{tcdA}$*::mNG*) grown to stationary phase in TY broth. The mNeonGreen signal for the $P_{tcdA}$*::mNG* reporter in Δ*rstA** was decreased (marked as ADJUSTED) so that differences in $P_{tcdA}$*::mNG* reporter expression between WT* and Δ*tcdR** could be visualized. Scale bar = 5 µm. **b–c.** Superplots of single-cell fluorescence intensity for $P_{cwp2}$*::LP-mScI3.* (**b**) and $P_{tcdA}$*::mNG* (**c**) quantified using SuperSegger[56] from the strains shown in **a**. The outlined colored dot represents the median fluorescence measured for a given biological replicate. The horizontal black line indicates the mean fluorescence determined for three biological replicates. The dotted red line in **b** marks the fluorescence cutoff for $P_{cwp2}$*::LP-mScI3*, while the dotted red line in **c** is used to define Toxin-ON in panel **d** (n = 200 cells per condition). **d.** Percentage of cells expressing the toxin-specific $P_{tcdA}$*::mNG* reporter in the indicated dual reporter strains. % Toxin-ON was determined as the proportion of cells with mNeonGreen signal greater than one standard deviation above the mean fluorescence of the Δ*tcdR* dual reporter strain. The median value for a given biological replicate based on analyses of 200 cells was determined (colored outlined dots) and then averaged to give the average % Toxin-ON. Mean and standard deviation are shown for three biological replicates (n = 600 cells). Statistical significance was determined using a one-way ANOVA and Tukey's test. Only statistically significant comparisons are shown.

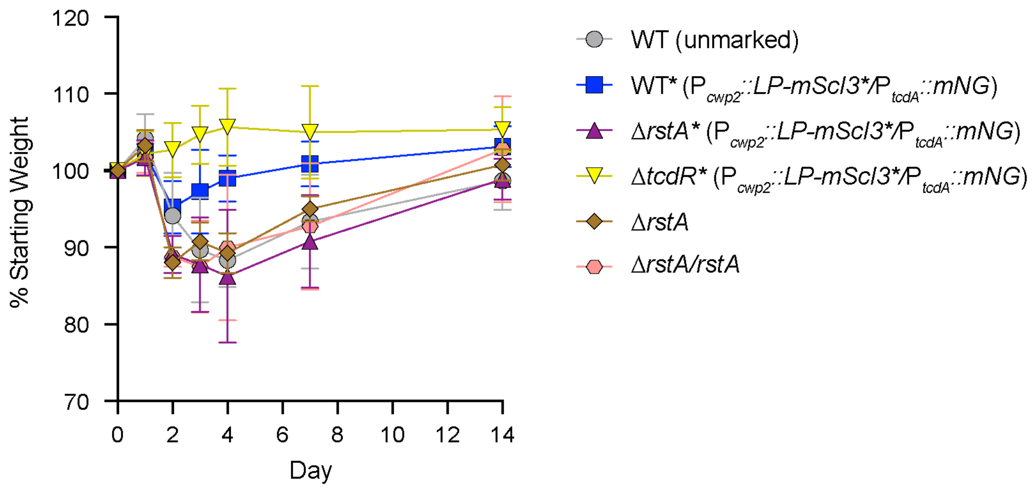


**Extended Data Figure 2. Percent weight change of mice over the course of infection with the indicated dual reporter or unmarked strains**. Mice were infected with 1 × $10^6$ spores of the indicated *C. difficile* strains. Values are relative to baseline weight at the time of infection (WT, n = 23; WT*, n = 8; Δ*rstA**, n = 11; Δ*tcdR**, n = 4; Δ*rstA*, n = 4; Δ*rstA* /*rstA*, n = 4, * = $P_{cwp2}$*::LP-mScI3*/$P_{tcdA}$*::mNG*).

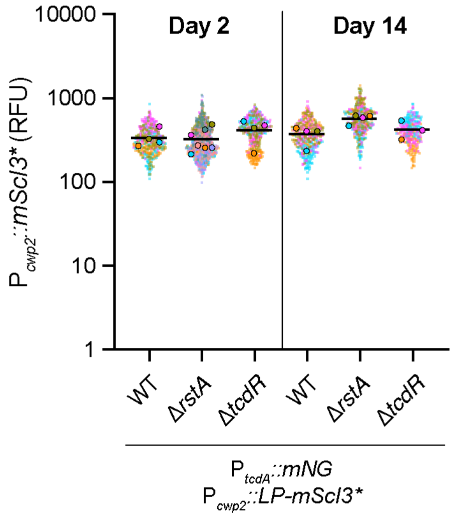


**Extended Data Figure 3. Quantification of constitutive reporter fluorescence during murine infection.** Superplot of constitutive reporter fluorescence ($P_{cwp2}$*::LP-mScI3*) at the single-cell level in colonic sections of mice infected with the indicated dual reporter strains Day 2 (48 hours) or 14 days post-infection. The median value of 100 cells per mouse was determined (colored outlined dot) and then averaged to give the mean fluorescent value (horizontal black line) (n = 400 cells per condition; Δ*rstA* at Day 2, n = 700).

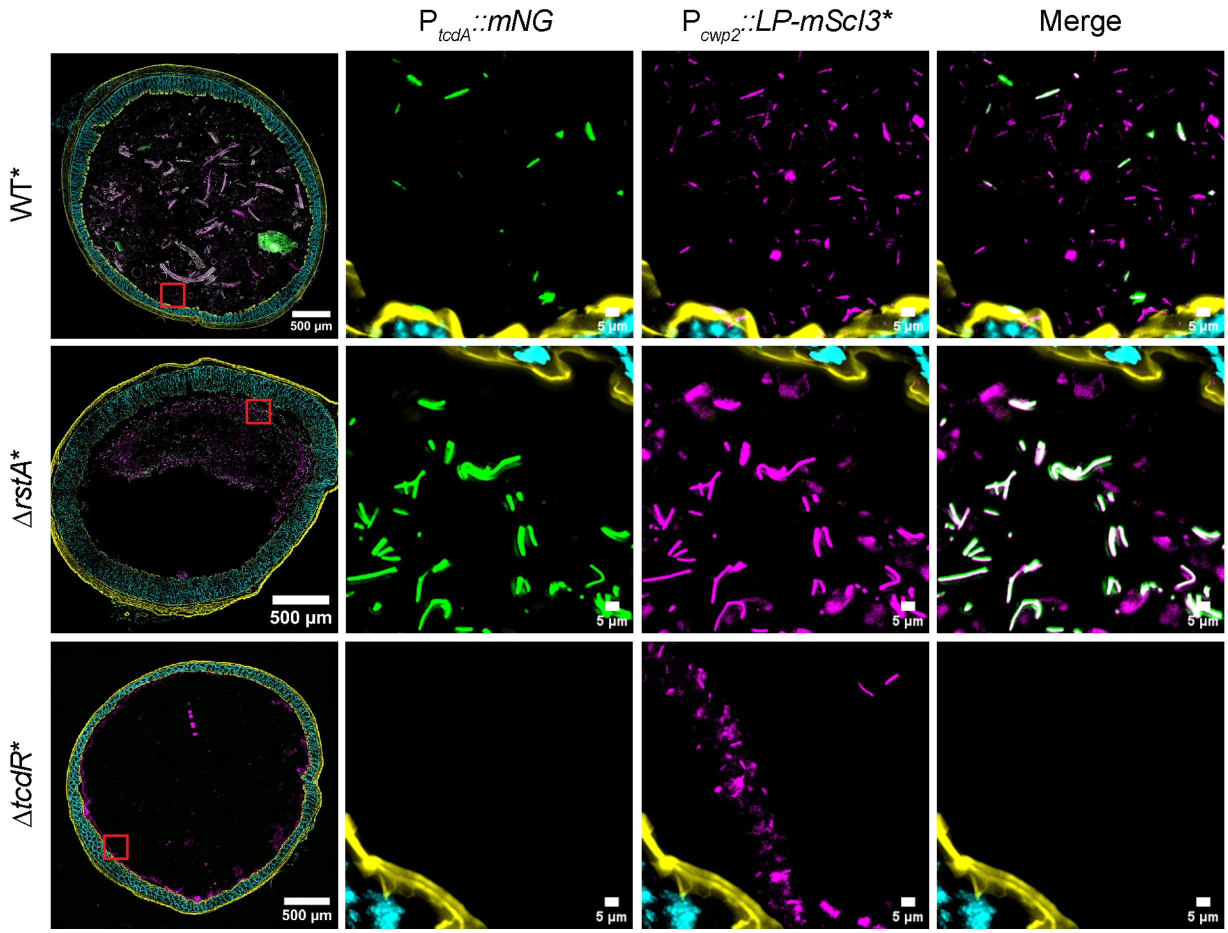


**Extended Data Figure 4. Representative images of colonic sections harvested from mice for dual reporter strains**. Mice were infected with WT*, Δ*rstA**, or Δ*tcdR** dual reporter strains (* = P$_{cwp2}$::*LP-mScI3* / P$_{tcdA}$::*mNG*) for 48 hrs before colon tissues were harvested, fixed in 4% PFA for 4 hours, cross-sectioned, and embedded in OCT. 10 µm sections were stained with phalloidin (F-actin, yellow) and DAPI (nuclei, cyan). Inset region is boxed in red.

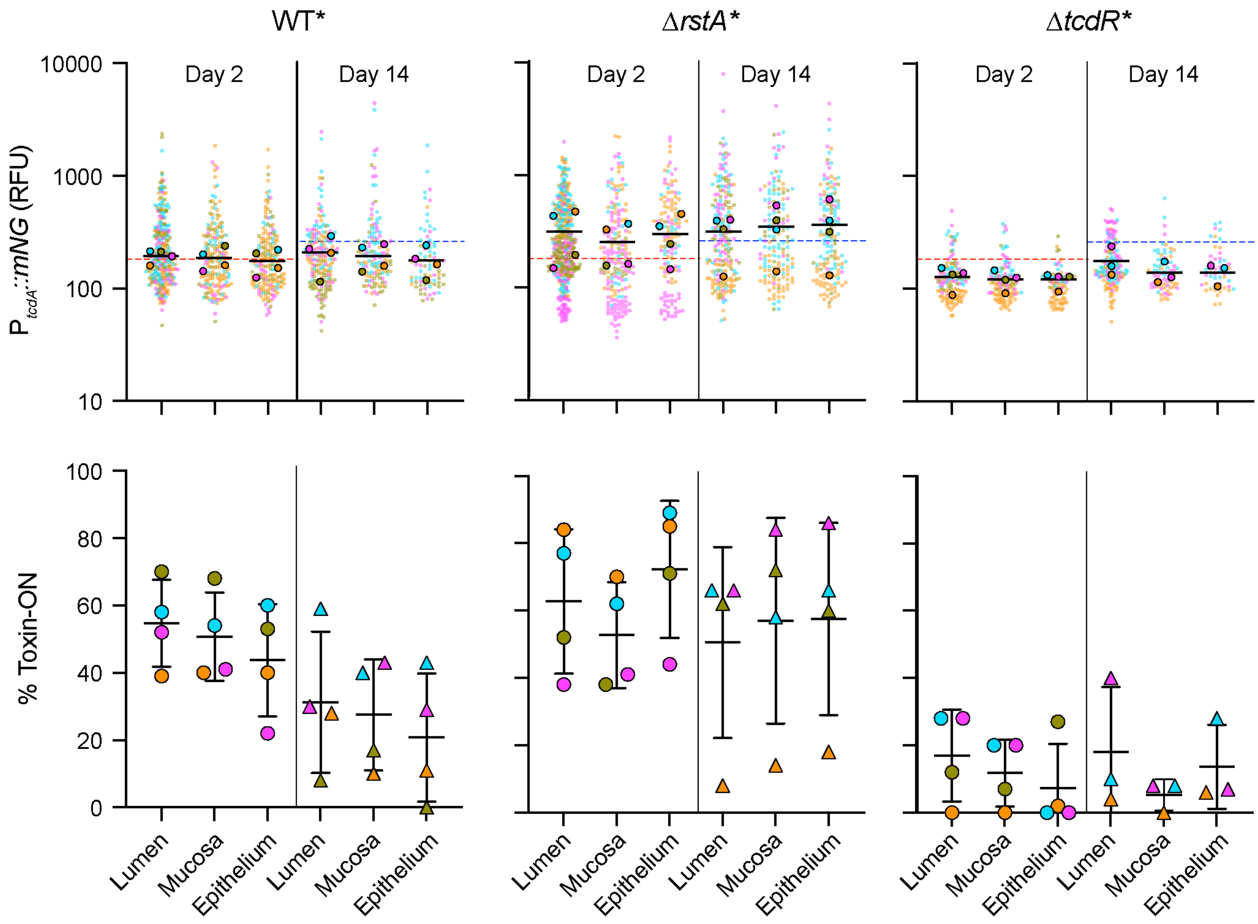


**Extended Data Figure 5. Spatial distribution of toxin gene expression during murine infection. a.** Superplot of toxin gene-specific reporter fluorescence ($P_{tcdA}$::*mNG*) at the single-cell level in colonic sections of mice infected with the indicated dual reporter strains (* = $P_{cwp2}$::*LP-mScI3*/$P_{tcdA}$::*mNG*) at Day 2 or Day 14 post-infection. Cells were localized in the lumen, mucosa, or epithelium by visualizing only in the mScI3 channel to avoid bias toward high $P_{tcdA}$::*mNG* expression. Colored outlined dots represent the median value of counted cells per mouse per time point and location in the colon (lumen, mucosa, epithelium). The horizontal black line indicates the average of the median fluorescence values determined for each mouse. For WT* at Day 2, a total of 300 cells were counted in the lumen (Mouse A, n = 100; B, n = 100; C, n = 50; D n = 50); 187 cells were counted in the mucus (A, n = 100, B, n = 37; C, n = 22; D, n = 28). A total of 183 cells were counted in the epithelium (A, n = 100; B, n = 20; C, n = 23; D, n = 40). For Δ*rstA** at Day 2, a total of 400 cells were counted in the lumen (A, n = 100; B, n = 100,; C, n = 100; D, n = 100). A total of 185 cells were counted in the mucus (A, n = 40; B, n = 21; C, n = 100; D, n = 24). A total of 134 cells were counted in the epithelium (A, n = 53; B, n =

19; C, n = 55; D, n = 7). For Δ*tcdR** at Day 2, a total of 125 cells were counted in the lumen (A, n = 50; B, n = 25; C, n = 25; D, n = 25). A total of 114 cells were counted in the mucus (A, n = 50; B, n = 25; C, n = 25; D, n = 14). A total of 87 cells were counted in the epithelium (A, n = 50; B, n = 15; C, n = 11; D n = 11).

For WT* at Day 14, a total of 189 cells were counted in the lumen (A, n = 50; B, n = 39; C, n = 50; D, n = 50). A total of 139 cells were counted in the mucus (A, n = 30; B, n =43; C, n = 37; D n = 29). A total of 81 cells were counted in the epithelium (A, n = 9; B, n = 28; C, n = 17; D, n = 27). For Δ*rstA** at Day 14, a total of 200 cells were counted in the lumen (A, n = 50; B, n = 50; C, n = 50; D, n = 50). A total of 150 cells were counted in the mucus (A, n = 50; B, n = 50; C, n = 25; D, n = 25). A total of 147 cells were counted in the epithelium (A, n = 50; B, n = 50; C, n = 22; D, n = 25). For Δ*tcdR** at Day 14, a total of 150 cells were counted in the lumen (A, n = 50; B, n = 50; C, n = 50). A total of 75 cells were counted in the mucus (A, n = 25; B, n = 25; C, n = 25). A total of 50 cells were counted in the epithelium (A, n = 17; B, n = 18; C, n = 15). **b.** Percentage of cells expressing the toxin-specific $P_{tcdA}$*::mNG* reporter in the indicated dual reporter strains in colonic sections in either the lumen, mucosa, or epithelium from mice 2 or 14 days post-infection. % Toxin-ON was determined as the proportion of cells with mNeonGreen signal greater than one standard deviation above the mean fluorescence of the Δ*tcdR** dual reporter strain (dashed red line). Mean and standard deviation are shown based on percentages measured from four mice (cells quantified per mouse stated above).

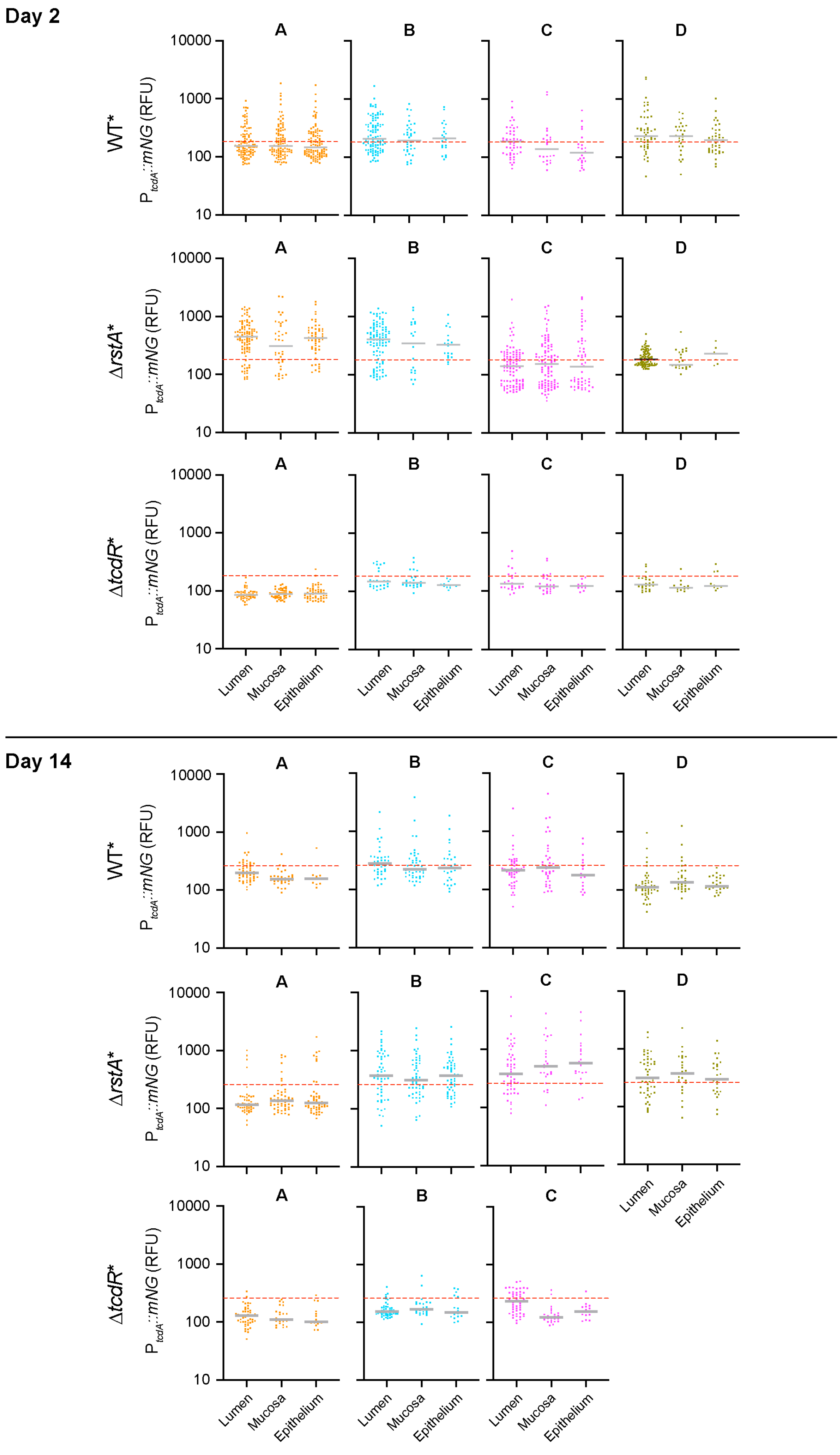


78 **Extended Data**

**Figure 6. Spatial distribution of toxin gene expression in individual mice.** Plots of toxin-specific reporter fluorescence ($P_{tcdA}$*::mNG*) at the single-cell level in colonic sections of mice infected with the indicated dual reporter strains 2 or 14 days post-infection. Cells were localized in the lumen, mucosa, or epithelium by visualizing only in the mScI3 channel to avoid bias toward high $P_{tcdA}$*::mNG* expression. Each letter (A–D) represents a different mouse colonized with the indicated strain. Colored dots represent the median single-cell $P_{tcdA}$*::mNG* fluorescence measured for a given mouse. The median fluorescence values were averaged to determine the geometric mean fluorescence per location. The dotted red line indicates the cutoff for Toxin-ON cells, defined as one standard deviation above the mean fluorescence of the Δ*tcdR* dual reporter strain at the indicated time point (For cell counts per mouse per location, see previous section *Extended Data Figure 5. Spatial distribution of toxin gene expression during murine infection*).

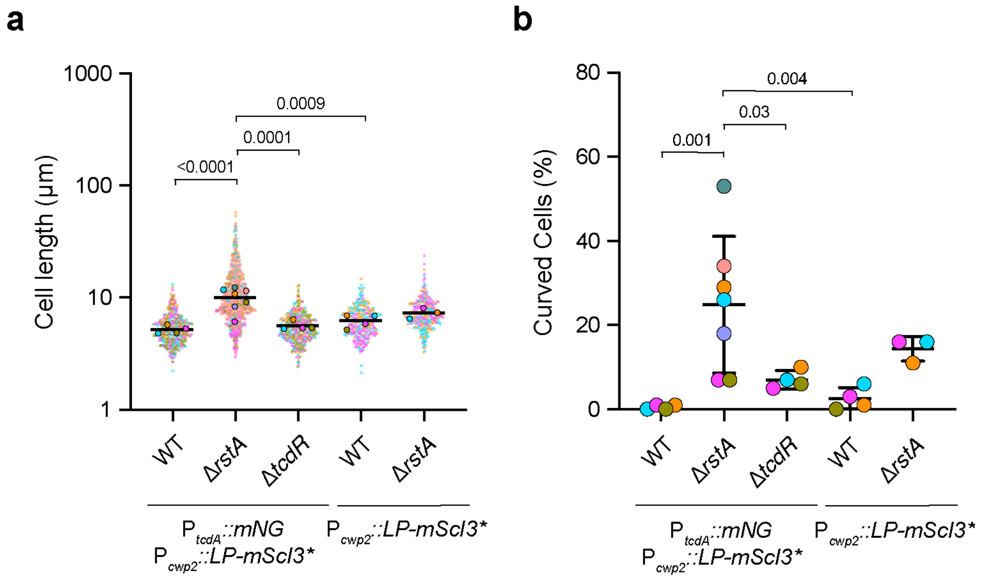


**Extended Data Figure 7**. **Comparison of the cell morphology of dual vs. single reporter strains during infection. a.** Superplots of cell lengths measured for individual cells in colonic sections for dual reporter strains in the WT, Δ*rstA*, and Δ*tcdR* backgrounds, as well as single constitutive $P_{cwp2}$*::LP-mScI3*-only strains in the WT and Δ*rstA* backgrounds. **b.** Percentage of curved cells for the dual reporter in WT, Δ*rstA*, and Δ*tcdR* backgrounds, as well as single constitutive $P_{cwp2}$*::LP-mScI3*-only strains in the WT and Δ*rstA* backgrounds. For both plots, the median value of 100 cells per mouse was determined (colored outlined dot) and then averaged to give the mean fluorescent value (horizontal grey line) (n = 400 cells per condition; Δ*rstA*($P_{cwp2}$*::LP-mScI3*/$P_{tcdA}$*::mNG*) at Day 2, n = 700). Lines show the geometric mean and standard deviation based on measurements made per mouse. Statistical significance was determined using a one-way ANOVA and Tukey's test. Only statistically significant comparisons are shown.

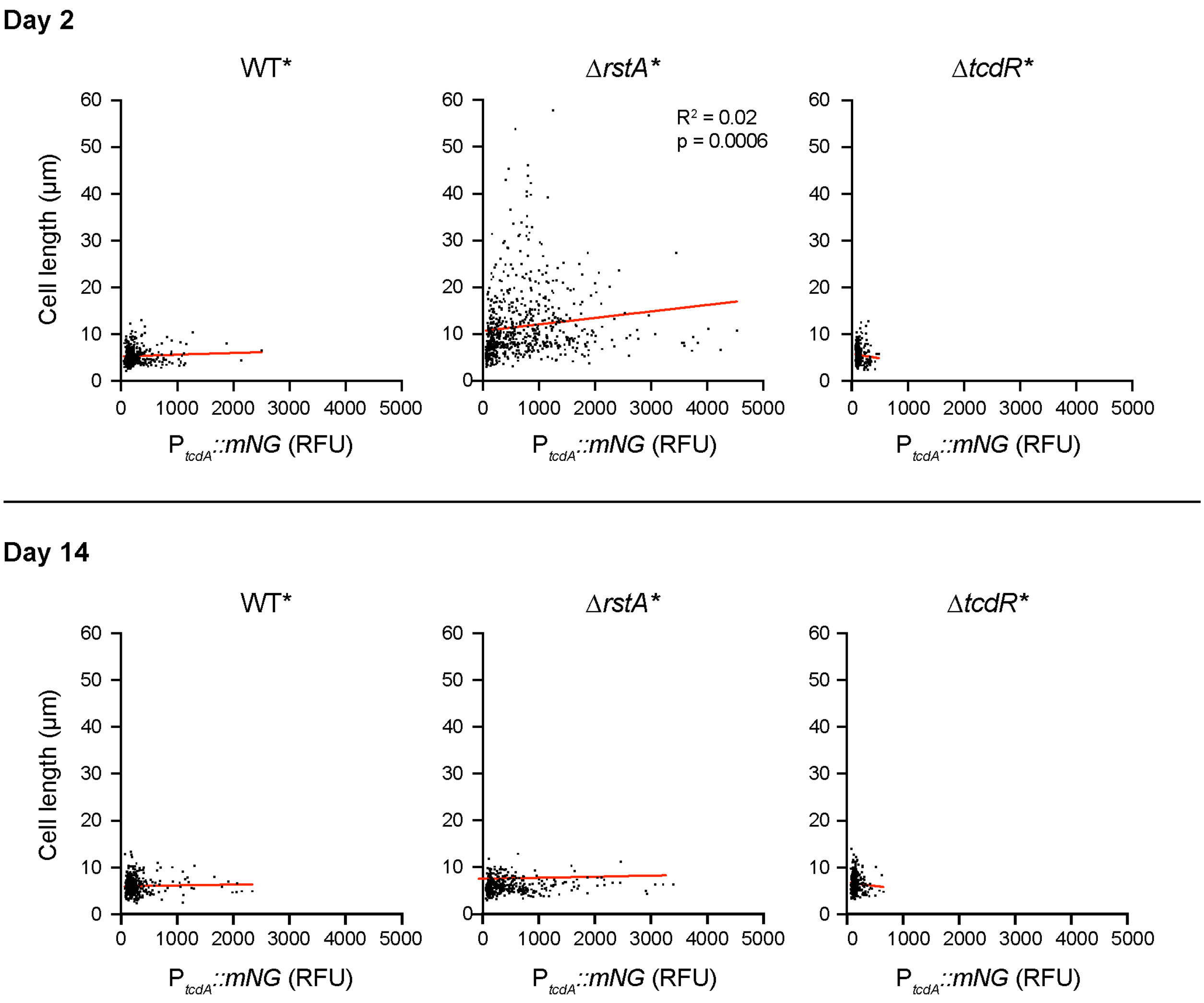


**Extended Data Figure 8. Relationship between toxin gene expression and cell length during murine infection**. Scatter plots showing $P_{tcdA}$::*mNG* fluorescence in WT*, Δ*rstA**, and Δ*tcdR** dual reporter strains (* = $P_{cwp2}$::*LP-mScI3*/$P_{tcdA}$::*mNG*) at Day 2 and Day 14 post-infection vs. individual *C. difficile* cell length (µm). Linear regression lines (red) are shown for each strain and timepoint (n = 4 mice, 100 cells per mouse, 400 total cells per condition; Δ*rstA* at Day 2, n = 7 mice, 700 total cells). The only significant correlation observed was for the Δ*rstA* dual reporter strain at Day 2.

114

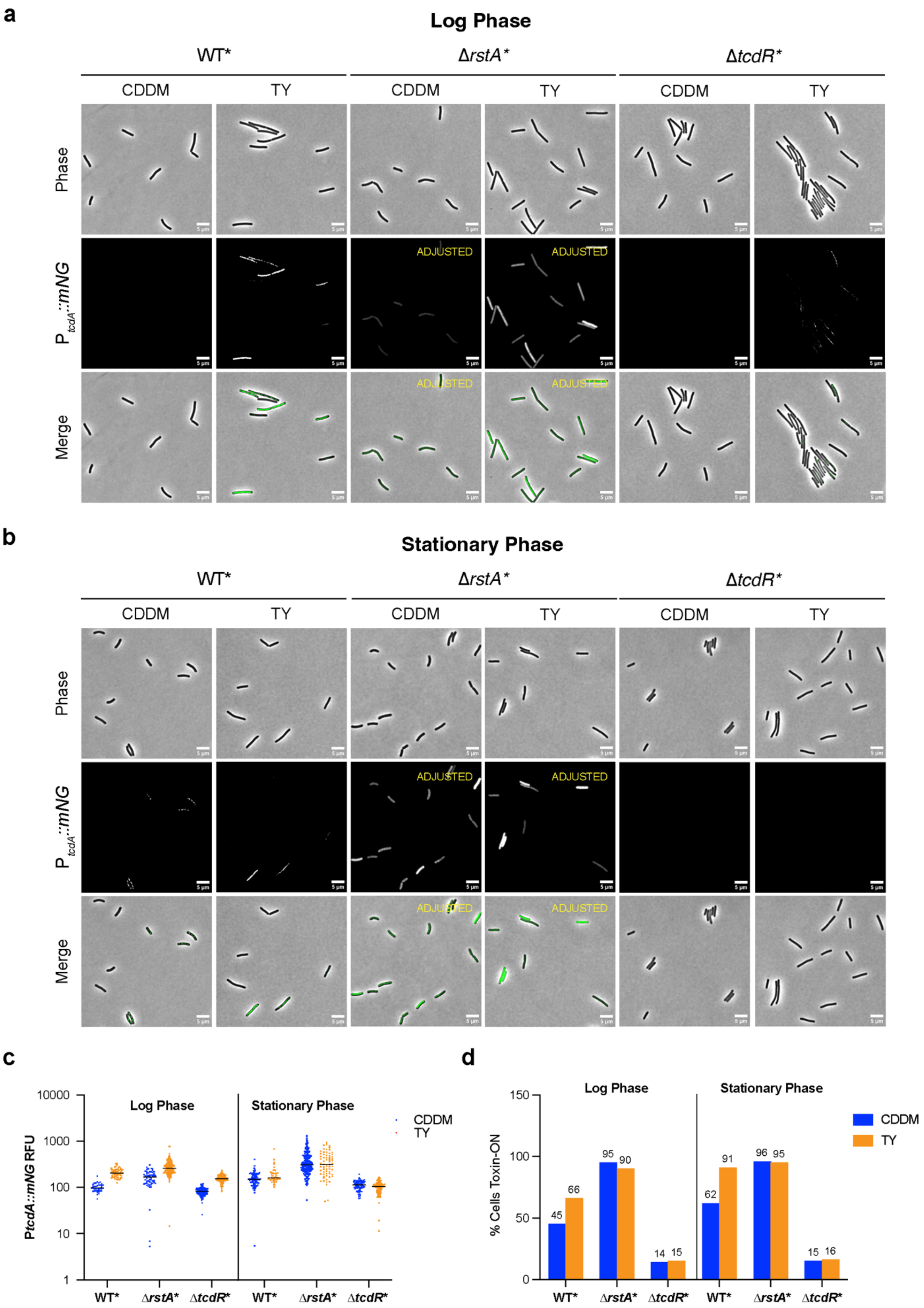
a
Log Phase
WT*
ΔrstA*
ΔtcdR*
CDDM
TY
Phase
PtcdA::mNG
Merge
ADJUSTED
b
Stationary Phase
c
PtcdA::mNG RFU
10000
1000
100
10
1
Log Phase
Stationary Phase
CDDM
TY
d
% Cells Toxin-ON
150
100
50
0
45
66
95
90
14
15
62
91
96
95
15
16

**Extended Data Figure 9. Toxin gene expression in dual reporter strains during growth in nutrient-rich TY or minimal CDDM broth at mid-log or stationary phase**. **a-b.** Representative fluorescence microscopy of the indicated dual reporter strains grown in either nutrient-rich TY or minimal CDDM broth at mid-log or stationary phase. **c.** Single-cell quantification of $P_{tcdA}$::*mNG* signal from images in **a** and **b**. Each dot represents the median mNG fluorescence of an individual cell. The black line represents the median across all single-cell median values. **d.** Percentage of cells expressing the toxin-specific $P_{tcdA}$::*mNG* reporter in the indicated dual reporter strains grown in TY or CDDM, at either log-phase or stationary phase. % Toxin-ON was defined as the proportion of cells with mNeonGreen fluorescence greater than one standard deviation above the mean fluorescence of the Δ*tcdR* dual reporter strain under the same media and timepoint conditions.
